# Uptake and determinants of cervical cancer prevention services among female college students in Kenya: A cross-sectional survey

**DOI:** 10.1101/2020.05.04.076513

**Authors:** Elisha Ngetich, Irene Nzisa, Alfred Osoti

## Abstract

**Introduction:** Cervical cancer is the leading cause of cancer death in women in low- and middle-income countries. In Kenya, cervical cancer incidence and prevalence have been increasing and in 2018 alone, there were 3286 deaths from cervical cancer. Previously, studies on cervical cancer prevention strategies have focused on women above 30 years old. However, as the risk factors for cervical cancer are acquired as early as in the teen years, an understanding of the awareness, uptake and determinants of screening services among college female students will help inform prevention strategies. This study sought to determine the awareness, uptake, determinants and barriers to cervical cancer screening services among colleges students in Kenya.

**Methods:** This was a multicenter cross-sectional study conducted in eight universities spread all over Kenya. Participants were interviewed using a self-administered structured questionnaire on sociodemographics, reproductive history, awareness on cervical cancer including screening practices, and attitude towards cervical cancer prevention services. Descriptive statistics were summarized using means and standard deviation (SD) for parametric data and median and interquartile range (IQR) for non-parametric data. Univariable and multivariable logistic regression analyses were done to determine odds ratios of factors associated with uptake of cervical cancer screening services. P-value of <0.05 was considered statistically significant.

**Results:** Between January 2017 and Sept 2017, we screened 800 and enrolled 600 female colleges students from eight universities in Kenya. In total, 549 of the 600 (92%) participants completed the questionnaire. The median age (IQR) was 21(20,22) years. Nearly two-thirds 338(62.7%) were sexually active, while 54(16%) had concurrent sexual partners. The main form of contraception was oral postcoital emergency pills 123(64.7%). Only 76(14.4%) had screened for cervical cancer, and the commonest approach was a Pap smear 47(61.8%). About one half 40(54.1%) did not like their experience due to pain, discomfort and bleeding. Four out of five participants (439, 80.7%) had poor knowledge on cervical cancer screening. On bivariate analysis, increased level of awareness (odds ratio [OR] 1.08 95% Confidence Interval [CI] 1.03,1.18, p = 0.004), knowledge of someone with cervical cancer(OR 0.43 CI 0.23,0.78 p=0.006) and a perception of self-risk (OR2.6 CI 1.38,4.98 p=0.003) were associated with increased odds of uptake of cervical cancer screening. In the multivariate analysis, high awareness was significantly associated with increased odds of cervical cancer screening (OR 1.12 CI 1.04, 1.20 p=0.002).

**Conclusions:** Female college students in Kenya had low levels of awareness on cervical cancer and had very low uptake of cervical cancer screening. However, high perception of self-risk and perceived benefit was associated with increased odds of cervical cancer screening.

**Recommendations:** Since female colleges students are generally thought to be more knowledgeable and have better access to information compared to the general population, the low levels of knowledge and uptake of cervical cancer screening, calls for a rethink of strategies that focus on the younger population including those in primary, high school and universities. Such strategies include HPV vaccination and incorporation of cancer prevention in school curriculum.

## INTRODUCTION

Cervical cancer is the second leading cause of cancer deaths in low and middle-income countries LMICs (1). The World Health Organization (WHO) estimates that in 2018 there were 570,000 new cases and 310,000 deaths globally from cervical cancer. Most, (85%) of these deaths were from LMICs(2). Despite increased focus on cervical cancer prevention and treatment, LMICs still lack well-structured control programs including screening and treatment of precancerous lesions(3,4). In Kenya, cervical cancer is the leading cause of cancer mortality among women. In 2018, Kenya had 5,250 new cases and 3, 286 deaths from cervical cancer, and 40% of women with normal cytology had high risk Human Papilloma Virus (HPV)(5), the cause of cervical cancer.

Sub-Saharan Africa (SSA) has the highest burden of cervical cancer owing to high prevalence of its risk factors including the close link between Human Immunodeficiency Virus HIV and HPV (6–8). Notably, Antiretroviral Therapy (ART) has been associated with reduced risk of HPV, precancerous lesions and regression of Cervical Intraepithelial Neoplasia (CIN)2+ lesions(9). Other risk factors common in SSA include early sexual debut, multiple sexual partners,, smoking, multiparity (10–14). In addition, high poverty rates reduce access to screening, early diagnosis and prompt management of precancerous lesions in SAA despite improved approaches to screening and management globally(15,16).

Effective screening approaches including cytology-based screening like Pap Smear may not be sustainable for most of the African countries because of the cost and time it takes to have the results(17). This makes affordable techniques that provide instant results like Visual Inspection with Acetic Acid (VIA) and Visual Inspection with Lugol’s Iodine (VILI) sustainable alternatives (18,19). However, such interventions are only effective if delivered to the target population. Although, the progression of cervical cancer from HPV infection to invasive carcinoma can take about 20 years, thus providing adequate time for intervention(20), HIV increases this rate significantly, hence the need for prompt intervention among women living with HIV/AIDS(21). Furthermore, since HPV infection is acquired soon after sexual debut, targeting much younger population especially in high school and university may be necessary(22). Thus, cervical cancer awareness and screening remains a priority.

Despite tremendous improvements in screening, prevention and management approaches, the incidence and prevalence of cervical cancer continues to increase in LMICs, including Kenya. To better understand the reasons for the increasing burden for cervical cancer in Kenya, an assessment of the current uptake of the screening services and level of awareness on cervical cancer particularly among at risk population prior to onset of disease is necessary. This study sought to evaluate the level and determinants of awareness and uptake of cervical cancer screening among female college students in Kenya, so as to identify gaps for potential intervention.

## METHODS

### Study design

This was a multi-center cross-sectional study conducted from January 2017 to September 2017 among college students in Kenya.

### Study setting

This study was conducted in eight institutions of higher learning spread all across Kenya, namely University of Nairobi, Kenyatta University, Moi University, Jomo Kenyatta University of Science and Technology, Daystar University, Pumwani college and Technical University. Participating institutions were selected using multi-stage random sampling.

### Study participants

Potential participants, female college students, were identified from different departments and all academic years using multi-stage random sampling. All potential participants received detailed explanation of study procedures and were asked to provide consent prior to enrolment. The study was approved by the University of Nairobi/KNH Ethical Review Committee. All participants signed written informed consent.

### Sample size

We estimated that a sample size of 600 participants proportionally divided among the universities would detect 20% uptake of cervical cancer screening among the female college students.

### Study procedures

#### Data collection and tools

Consenting participants were interviewed using self-administered paper questionnaires under the supervision of research assistants. The study questionnaire had five components on sociodemographic, obstetric and gynecological history, awareness of cervical cancer, cervical cancer screening practices, and barriers and attitude towards cervical cancer screening. The 30-item standardized questionnaire that had mainly close-ended questions was reviewed by two independent public health experts and pretested before data collection. Questionnaires were serially numbered and there were no participant identifiers to ensure confidentiality.

#### Outcome

The main outcome was self-reported uptake of cervical cancer screening in which participants were asked if they had ever had cervical cancer screening and if so, the method used. Secondary outcomes were level of awareness on cervical cancer and barriers and attitudes towards uptake of cervical cancer screening.

#### Data analysis

Data from all the eight centers were entered into excel sheet and collated and exported to STATA^®^ version *14*. College Station, TX: StataCorp LP for analysis. To assess the level of awareness on cervical cancer questions on risk factors, symptoms, diagnosis and treatment of cervical cancer were scored depending on the number of correct responses. The maximum possible score was 26 and the minimum zero. Scores below 14 were classified as poor and 14 or higher as good knowledge.

Descriptive statistics were summarized for all the baseline characteristics using means and standard deviations for parametric, median and quartile ranges for non-parametric and proportions for categorical variables. We used Student t-tests and nonparametric Wilcoxon rank sum tests versus Pearson’s chi-square tests or Fisher’s exact tests to compare continuous variables and categorical variables. We obtained crude and adjusted odds ratio estimates of the association between cervical cancer awareness score and screening practice using logistic regression model in bivariate and multivariable analysis adjusted for year of study, perceived risk and benefit, as potential confounding variables. In multivariable logistic regression we also accounted for participant year of study and the courses. P-values <0.05 were considered statistically significant.

## RESULTS

Between January 2017 and September 2017, 800 Kenyan female colleges students from eight different universities spread across the entire country were approached and screened for eligibility and 600 enrolled. In total 549 (92%) participants completed the questionnaire. The baseline sociodemographic characteristics are described in **Table 1**. The median (IQR) age of the participants was 21 (20,22) years. **Figure 2** shows a box plot of the age of the participants. The youngest participant was 18 years while the oldest was 30 years old.

**Table 1:** Sociodemographic characteristics of Kenyan female college students interviewed on knowledge and uptake of cervical cancer screening (N= 549)

| **Characteristic** | **Frequency (n)** | **Percentage (%)** |
| --- | --- | --- |
| **University** |  |  |
| Daystar | 21 | 3.8 |
| Egerton | 29 | 5.3 |
| Jomo Kenyatta | 82 | 14.9 |
| Kenyatta | 125 | 22.8 |
| Moi | 50 | 9.1 |
| Pumwani | 13 | 2.4 |
| Technical | 50 | 9.1 |
| Nairobi | 179 | 32.6 |
| **Year of Study** |  |  |
| First | 61 | 11.1% |
| Second | 136 | 24.8% |
| Third | 205 | 37.3% |
| Fourth | 110 | 20.0% |
| Fifth y | 24 | 4.4% |
| Sixth | 13 | 2.4% |
| **Departments** |  |  |
| Agriculture and Veterinary sciences | 34 | 6.2% |
| Architecture and Engineering | 62 | 11.3% |
| Biological and Physical Sciences | 73 | 13.3% |
| Education and External Studies | 87 | 15.9% |
| Health Sciences | 90 | 16.4% |
| Humanities and social sciences | 203 | 37.0% |
| **Employment status** |  |  |
| Employed | 258 | 47.9 |
| Volunteer (n=258) | 75 | 28.3 |
| Salaried (n=258) | 190 | 71.7 |
| **Financial support** |  |  |
| Parents | 439 | 83.8 |
| Relatives | 38 | 7.3 |
| Self | 33 | 6.3 |
| Others | 14 | 2.7 |

### Sexual and reproductive history of female college students in Kenya

Participants’ sexual and reproductive history is summarized in **table 2**. About two-thirds (338, 63%) were sexually active. The media (IQR) age of sexual debut was 19(18,20). **Figure 2** represents a box plot of the age at sexual debut. One third (173, 34.1%) of those who were sexually active reported having more than one sexual partner and about one third (54, 31.2%) of whom were concurrent. In total, only about a third (188, 35.7%) of the study participants used contraceptives, the most common being emergency pills (123, 64.7%).

**Table 2:**
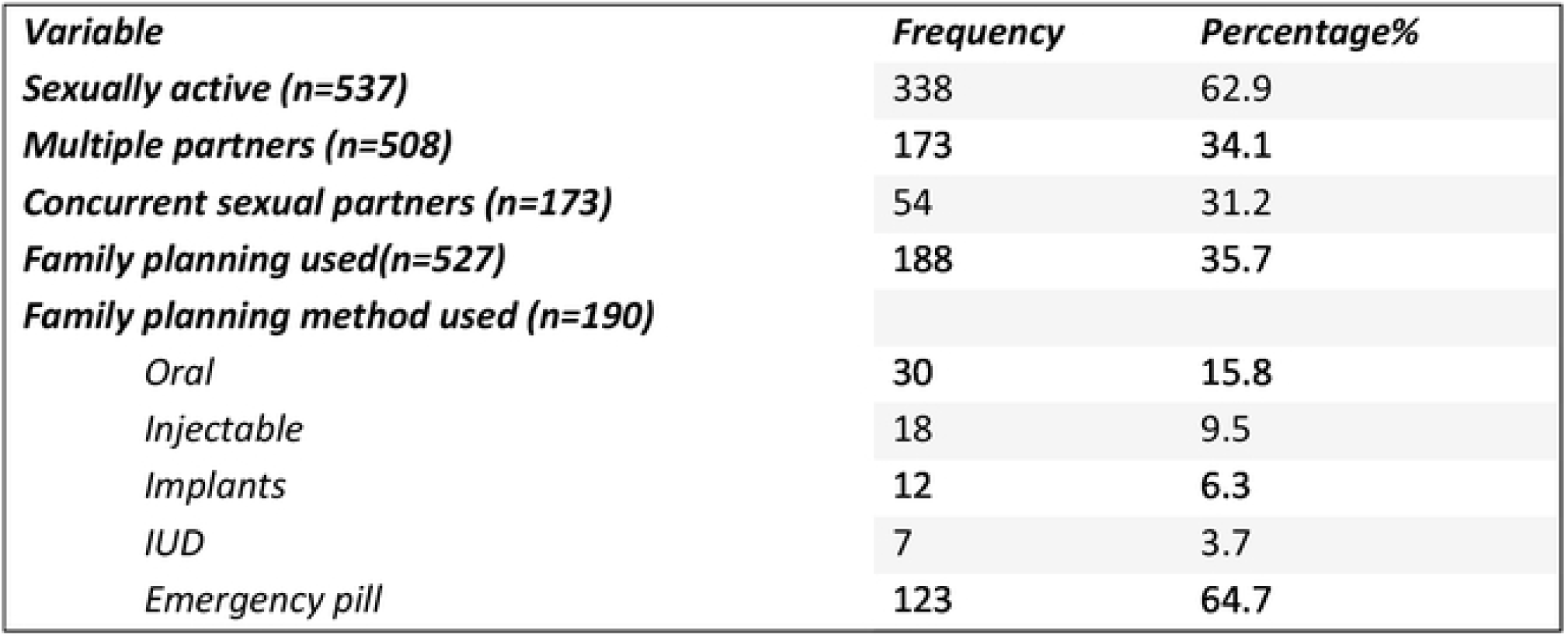
Sexual and reproductive history of Kenyan female college students interviewed on knowledge and uptake of cervical cancer screening (N=549)

### Level of awareness on risk factors, diagnosis and management of cervical cancer among female college students in Kenya

**Figure 1** is a bar chart of the level of awareness. Nearly all participants (538, 98%) of had heard of cervical cancer. The commonest source of information was social media (368,67.2%). Based on a score cut off of ≥13 out of 26, a paltry (105, 19.3%) had good knowledge on the risk factors, symptoms, diagnosis and treatment of cervical cancer. The mean (SD) score was 10 (±4).

**Figure 1:**
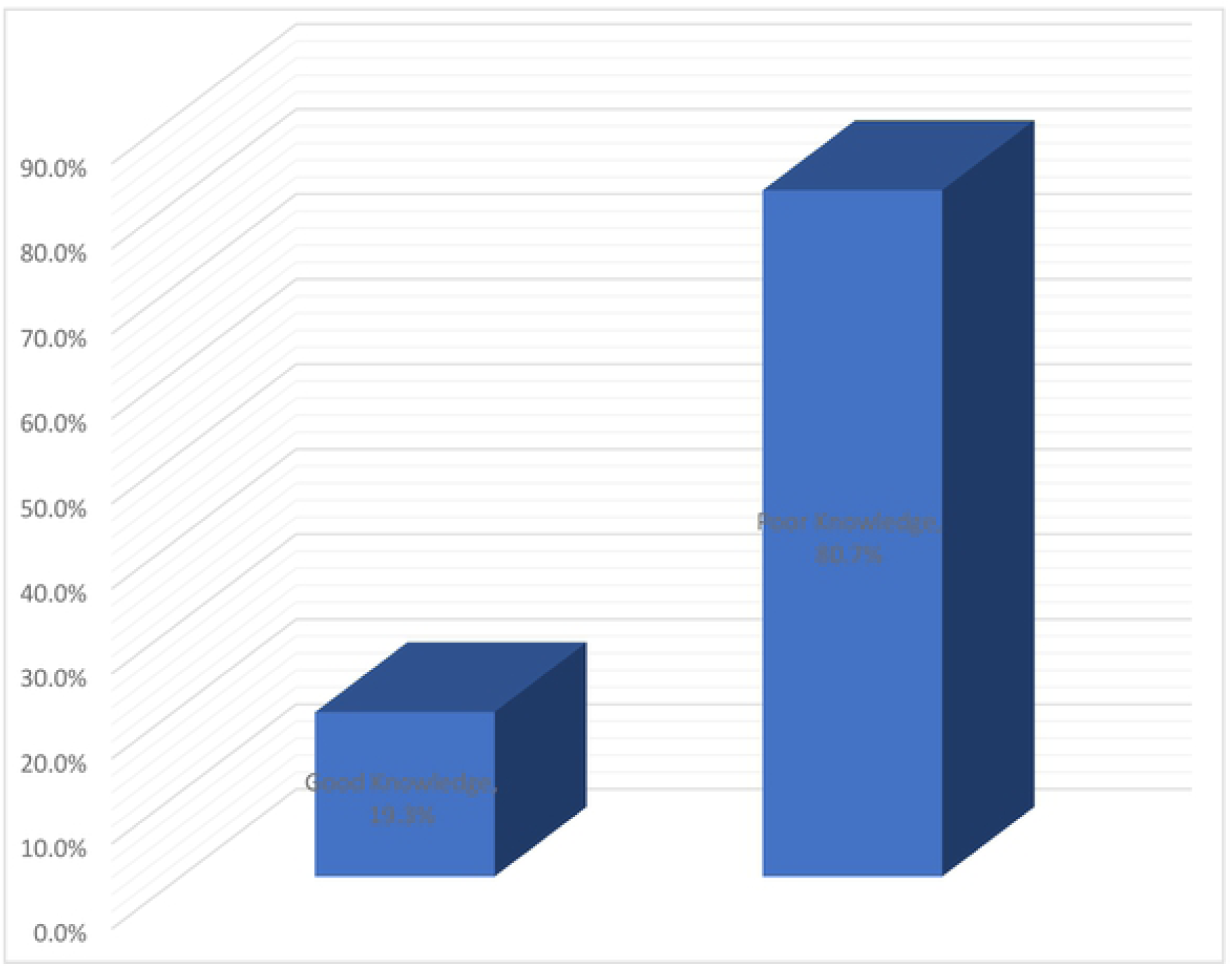
Level of awareness on risk factors, diagnosis and management of cervical cancer among Kenyan female college students.

**Figure 2:**
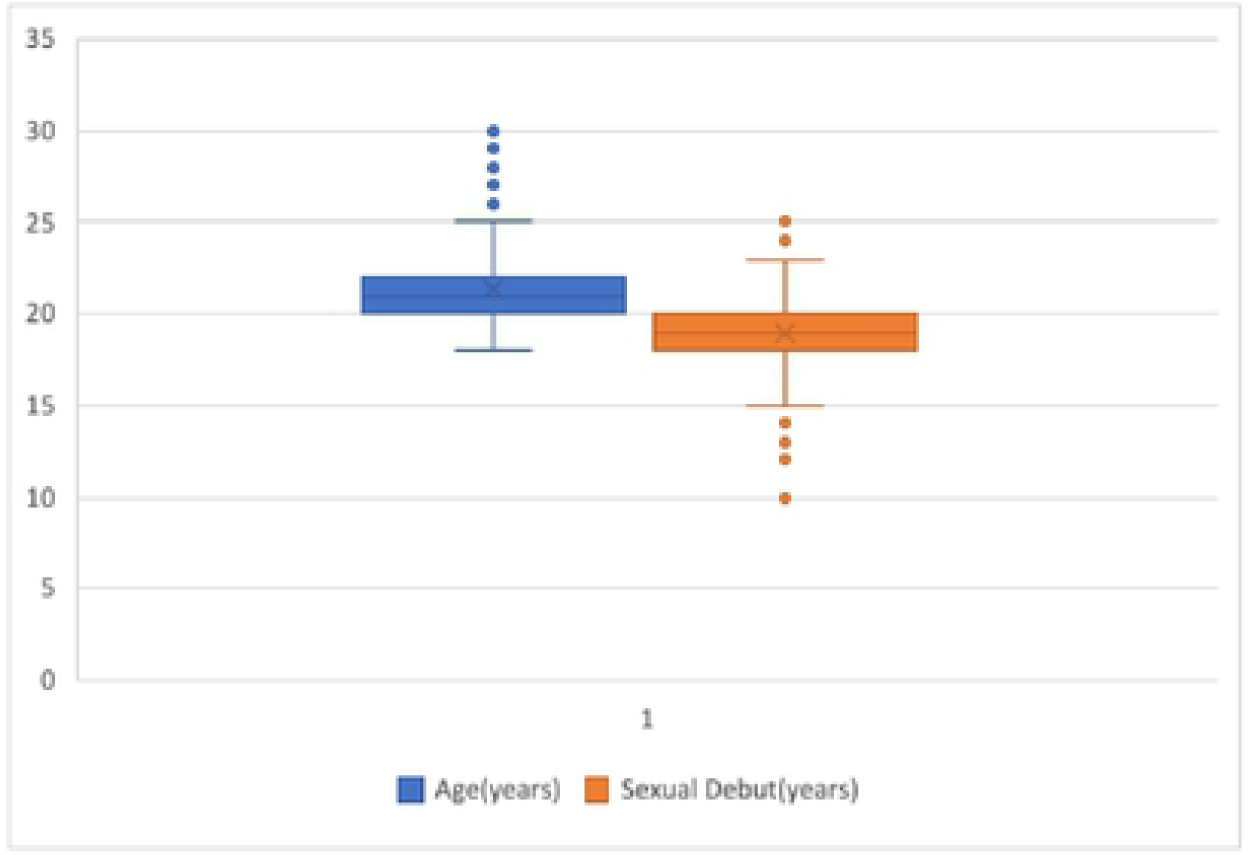
Box plot of participant age and age at sexual debut of Kenyan female college students interviewed on knowledge and uptake of cervical cancer screening

### Screening practices and acceptability of cervical cancer prevention services among female college students in Kenya

Screening practices and acceptability of cervical cancer prevention services are summarized in **table 3**. Of those who had heard of cervical cancer (338,98%) only (76,14.4%) had undergone cervical cancer screening. The most common screening methods were Pap smear (47,61.8%) followed by VIA (17,22.4%) then VILI (8,10.5%). HPV DNA testing was the least common (4,5.3%). Majority (73, 97%) of the screening results were normal with only one positive result. Notably, more than half (40,54.7%) of those who screened did not like their experience due to pain, discomfort and bleeding. About three-quarters (420, 76.5%) of the participants believed that screening for cervical cancer was beneficial and would recommend it to their friends and family. Reported perceived barriers to screening included lack of awareness on service availability, cost, discomfort and poor health care workers’ attitude.

**Table 3:** Cervical cancer screening practice and acceptability of prevention services among Kenyan female college students

| **Practice** | **Frequency** | **Percentage%** |
| --- | --- | --- |
| **Screening uptake (n=529)** |  |  |
| Ever screened | 76 | 14.4 |
| **Screening Location(n=76)** |  |  |
| Medical facility | 38 | 50 |
| Medical camp | 22 | 28.9 |
| College | 16 | 21.1 |
| **Screening findings(n=76)** |  |  |
| Normal | 73 | 97.3 |
| Abnormal | 1 | 1.3 |
| Other | 1 | 1.3 |
| TOTAL | 75 | 100.0 |
| **Experience(n=74)** |  |  |
| Liked it | 34 | 45.9 |
| Didn't like it | 40 | 54.1 |
| TOTAL | 74 | 100 |
| **Reasons for not liking it(n=38)** |  |  |
| Pain | 20 | 52.6 |
| Bleeding | 4 | 10.5 |
| Discomfort | 14 | 36.8 |
| TOTAL | 38 | 100.0 |
| **Screening Method(n=76)** |  |  |
| Pap smear | 47 | 61.8 |
| VIA | 17 | 22.4 |
| VILI | 8 | 10.5 |
| HPV testing | 4 | 5.3 |
| TOTAL | 76 | 100.0 |
| **Reasons not screened(n=266)** |  |  |
| Expensive | 61 | 22.9 |
| Uncomfortable | 101 | 38.0 |
| Perception of low self-risk | 104 | 39.1 |
| TOTAL | 266 | 100.0 |

In bivariate analysis, increased level of awareness was associated with increased odds of uptake of cervical cancer screening (Odds Ratio OR 1.083 Confidence Interval CI 1.03,1.18, p = 0.004), knowledge of someone with cervical cancer(OR 0.43 CI 0.23,0.78 p=0.006) and a perception of self-risk (OR2.6 CI 1.38,4.98 p=0.003. In the multivariate analysis, after adjusting for the year of study, knowledge of someone with cervical cancer, perceived benefit of screening and perceived self-risk, high awareness was associated with 12% greater odds of uptake of screening compared to those with low awareness (OR 1.12 CI 1.04, 1.20 p=0.002). **Table 4** shows the bivariate and multivariate logistic regression after adjusting for year of study, knowledge of someone with cervical cancer, perceived benefit of screening and perception of self-risk.

**Table 4:**
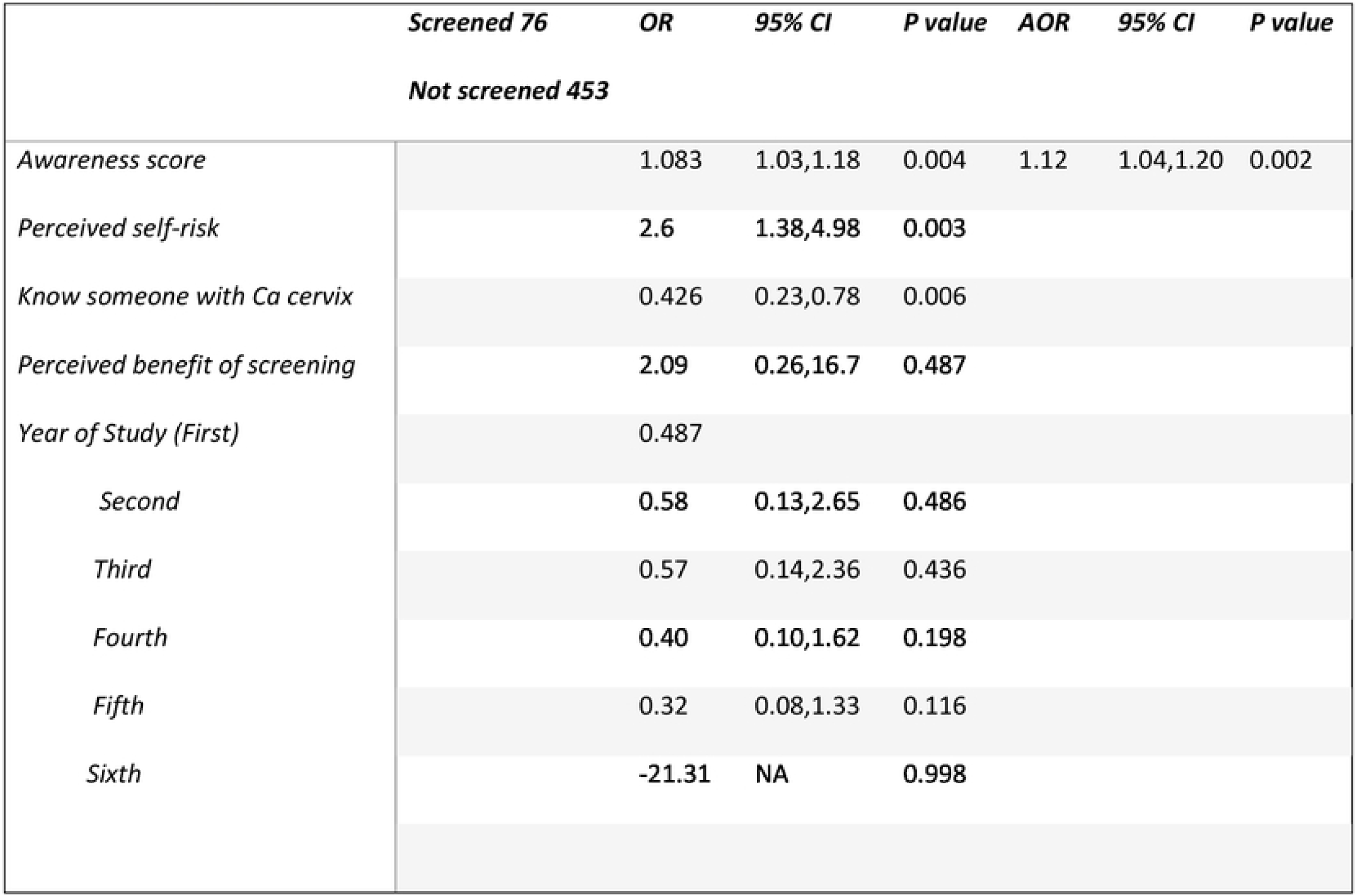
Association between cervical cancer awareness and uptake of screening services among Kenyan female college students

|  | *Screened 76* | *OR* | *95% CI* | *P value* | *AOR* | *95% CI* | *P value* |
| --- | --- | --- | --- | --- | --- | --- | --- |
|  | *Not screened 453* |  |  |  |  |  |  |
| *Awareness score* |  | 1.083 | 1.03,1.18 | 0.004 | 1.12 | 1.04,1.20 | 0.002 |
| *Perceived self-risk* |  | 2.6 | 1.38,4.98 | 0.003 |  |  |  |
| *Know someone with Ca cervix* |  | 0.426 | 0.23,0.78 | 0.006 |  |  |  |
| *Perceived benefit of screening* |  | 2.09 | 0.26,16.7 | 0.487 |  |  |  |
| *Year of Study (First)* |  | 0.487 |  |  |  |  |  |
| *Second* |  | 0.58 | 0.13,2.65 | 0.486 |  |  |  |
| *Third* |  | 0.57 | 0.14,2.36 | 0.436 |  |  |  |
| *Fourth* |  | 0.40 | 0.10,1.62 | 0.198 |  |  |  |
| *Fifth* |  | 0.32 | 0.08,1.33 | 0.116 |  |  |  |
| *Sixth* |  | -21.31 | NA | 0.998 |  |  |  |

## DISCUSSION

In this multicenter study representative of the Kenyan female college student population we found low uptake of screening services and poor knowledge on cervical cancer. Despite more than 98% having heard of cervical cancer and 62.9% reporting sexually activity, only 14.4% had undergone cervical cancer screening and over 80% scored poorly in the awareness score. This is worrisome considering that the chosen participants (university students) are perceived to be the most educated, socially empowered and with relatively better access to information compared to the rest of the population. It would therefore be expected that these participants should have a significantly higher awareness on cervical cancer. This is however not unique to Kenya. Similar findings of low uptake/ awareness of female college students were reported in India 0%, South Africa 9.8 %, Ghana 12%(23–25).

In our study, pain, discomfort and bleeding were reported as the commonest reasons for the poor uptake of cervical cancer screening. Some patients have previously described the procedure for pap smears as ‘scary’ as reported in the study by Gatumo et al(26). While patient reassurance and adequate relay of information before the procedure may help allay fear, these findings reiterate the need for better screening methods that are patient friendly, cost-effective and provide immediate results. Self-sampling and sending the sample for analysis is fast gaining preference globally and has been shown to increase women participation in cervical cancer screening(27,28). This is because it eliminates the use of speculums and unnecessary hospital visits for the procedure. However, self-sampling kits are fast gaining preference as they eliminate the uncomfortable use of speculums and unnecessary hospital visit for this procedure(27).

Evidently, this study population has little knowledge on cervical cancer and its prevention mechanisms. It is therefore not surprising that HPV vaccination programs and other cervical cancer prevention strategies including HPV vaccination would be met with reluctance and hesitancy(29). Onyebuchi et al(22) in his study on cancer prevention strategies in developing countries recommends a re-thinking of the sensitization strategies and presents a case for targeting teenagers and high school students as well as incorporating this into school curriculum. As Kenya and other LMICs of SSA continue with HPV vaccination program, this alone will not be effective in the absence of community sensitization and continued screening. Most importantly, however, is the recognition that even with advanced education, as demonstrated in this study, the decision to screen for cervical cancer may significantly be influenced by other factors including cultural and religious values that have a significant bearing on the decisions that people make on a daily basis(30–32).

The findings of this study should however be interpreted cautiously owing to inherent limitations. Data collection was self-reported and therefore may have been prone to recall and social desirability bias. Secondly, while the findings provide a lot of information on the need for more sensitization, they only apply to college level students and not lower level or out of school students. However, with universal access to primary and secondary education with nearly 100% transition to college the results can be widely applied. Finally, a complementary qualitative study would provide a better understanding of the reasons for the low uptake of screening services in this population.

In conclusion, awareness and uptake of cervical cancer screening services was shown to be poor among college students in this setting. While college students are thought to be most educated, socially empowered and with better access to information, this did not translate into better uptake of screening services and most still lead a risky sexual behavior with a significant number having multiple concurrent sexual partners. The findings from this study call for rethinking awareness creation on cervical cancer by targeting teenagers, high school and college students before they pick all the risk factors.

## Acknowledgement

We would likely to whole-heartedly thank the data collectors, supervisors and participants who dedicated their time and great effort in the data collection process.

